# Sporulation is dispensable for the vegetable-associated life cycle of the human pathogen *Bacillus cereus*

**DOI:** 10.1101/2021.01.15.426836

**Authors:** María Luisa Antequera-Gómez, Luis Díaz-Martínez, Juan Antonio Guadix, Ana María Sánchez-Tévar, Sara Sopeña-Torres, Jesús Hierrezuelo, Hung K. Doan, Johan H.J. Leveau, Antonio de Vicente, Diego Romero

## Abstract

*Bacillus cereus* is a common food-borne pathogen that is responsible for important outbreaks of food poisoning in humans. Diseases caused by *B. cereus* usually exhibit two major symptoms, emetic or diarrheic, depending on the toxins produced. It is assumed that after the ingestion of contaminated vegetables or processed food, spores of enterotoxigenic *B. cereus* reach the intestine, where they germinate and produce the enterotoxins that are responsible for food poisoning. In our study, we observed that sporulation is required for the survival of *B. cereus* in leaves but is dispensable in ready-to-eat vegetables, such as endives. We demonstrate that vegetative cells of *B. cereus* that are originally impaired in sporulation but not biofilm formation are able to reach the intestine and cause severe disorders in a murine model. We propose that loss of part of the sporulation programme and reinforcement of structural factors related to adhesion, biofilm formation and pathogenic interaction with the host are adaptive traits of *B. cereus* with a life cycle primarily related to human hosts. Furthermore, our findings emphasise that the number of food poisoning cases associated with *B. cereus* is underestimated and suggest the need to revise the detection protocols, which are based primarily on spores and toxins.

## Introduction

A major problem in food safety is the contamination of fresh and stored produce with human pathogens [1, 2], which may provoke food poisoning outbreaks [3]. *Bacillus cereus* is a natural inhabitant of soils and is able to live in association with plants. Members of this group of bacteria are also responsible for food poisoning outbreaks through consumption of raw vegetables and fruits or processed food [4, 5]. Food poisoning caused by *B. cereus* is classified into two main categories, emetic and diarrheic, depending on the battery of toxins produced [6]. An emetic poisoning is associated with the production of a dodecadepsipeptide called cereulide, [7, 8] a remarkably heat- and acid-stable toxin typically detected in high concentrations in processed food contaminated with *B. cereus* [9]. Diarrheic poisoning is known to occur after the ingestion of food contaminated with enterotoxigenic *B. cereus* spores which, upon germination in the intestine, produce heat-labile enterotoxins [10, 11]. A group of three primary enterotoxins are involved in diarrheic poisoning: haemolysin BL (Hbl), nonhaemolytic enterotoxin (Nhe) and cytotoxin K (CytK) [12, 13].

*B. cereus* spores are highly adhesive and resistant to stress, which ensures their persistence in a variety of environments, under various sanitation measures and in ready-to-eat or processed food [14]. Under more favourable environmental conditions, spores readily germinate to vegetative cells and begin new life cycles [15]. In addition to spores, contamination with *B. cereus* in the food industry is attributable to the formation of biofilms, in which bacterial communities assemble in response to a variety of signals that enable microorganisms to adapt to different environments [16]. In the proposed life cycle of *B. cereus*, these microbes live as saprophytes in soil, move to plants and later enter human-related environments (hospitals, the food industry or human bodies). Spores or vegetative cells rotate depending on the life cycle stage, and they are proposed to remain as individual cells or become embedded in biofilms.

In this study, we investigated the relevance of spores and biofilms of *B. cereus* to the persistence of this species on infrequently investigated plants. The comparative analysis of a group of isolates involved in food poisoning enabled us to propose that sporulation is a major ecological trait in the phylloplane (i.e. on leaf surfaces) but that sporulation is dispensable in ready-to-eat vegetables, where vegetative cells persist collectively within biofilms. Additionally, we demonstrate that intrinsic genetic defects in sporulation do not diminish the pathogenicity of vegetative cells of *B. cereus* in a murine model. In addition to these ecological and evolutionary implications, our study suggests that the number of food poisoning cases associated with *B. cereus* is underestimated, as the diagnosis of these cases is based primarily on spores and toxins.

## Materials and Methods

### Bacterial strains and growth conditions

Nine *B. cereus* isolates from food poisoning outbreaks were used in this study (**Table 1).** Bacteria were routinely precultured in LB broth (10 g/l tryptone, 5 g/l yeast extract and 5 g/l NaCl) or in 1.5% LB agar. *B. cereus* strains were grown overnight with shaking at 28°C. To study biofilm formation, *B. cereus* strains were cultured for 72 h at 28°C without shaking in TY broth (10 g/l tryptone, 5 g/l yeast extract, 5 g/l NaCl, 10 mM MgSO_4_ and 1 mM MnSO_4_) or 1.5% TY agar supplemented with 20 µg/ml and 10 µg/ml Congo Red and Coomassie Brilliant Blue filtered dyes, respectively [17]. For swarming motility assays, 0.7% TY agar was used [18]. For sporulation experiments, cells were cultured in Difco Sporulation Medium (DSM) [19] for 24 h with shaking at 28°C. To simulate *in vitro* gastric passage, the cells were incubated in J broth (JB) (5 g/l peptone, 15 g/l yeast extract, 3 g/l K_2_HPO_4_ and 2 g/l filtered glucose) [20].

**Table 1.**
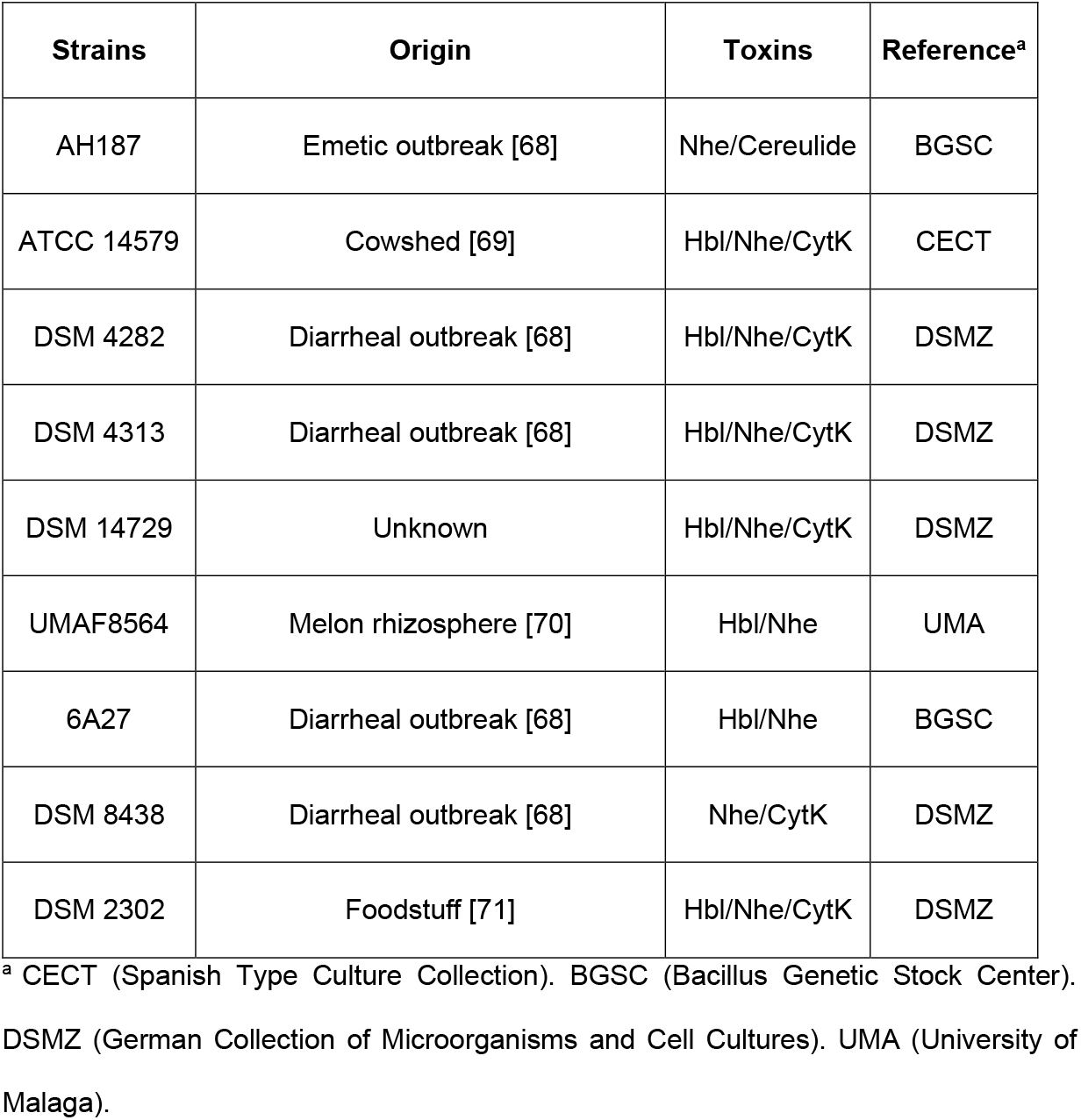
*Bacillus cereus* strains used in this study. Origin indicates the source where the strains were isolated. Toxins are the main enterotoxin or emetic toxin produced by each stain. Reference corresponds to the culture collection where the strains were obtained.

| Strains | Origin | Toxins | Reference<sup>a</sup> |
| --- | --- | --- | --- |
| AH187 | Emetic outbreak [68] | Nhe/Cereulide | BGSC |
| ATCC 14579 | Cowshed [69] | Hbl/Nhe/CytK | CECT |
| DSM 4282 | Diarrheal outbreak [68] | Hbl/Nhe/CytK | DSMZ |
| DSM 4313 | Diarrheal outbreak [68] | Hbl/Nhe/CytK | DSMZ |
| DSM 14729 | Unknown | Hbl/Nhe/CytK | DSMZ |
| UMAF8564 | Melon rhizosphere [70] | Hbl/Nhe | UMA |
| 6A27 | Diarrheal outbreak [68] | Hbl/Nhe | BGSC |
| DSM 8438 | Diarrheal outbreak [68] | Nhe/CytK | DSMZ |
| DSM 2302 | Foodstuff [71] | Hbl/Nhe/CytK | DSMZ |

<sup>a</sup> CECT (Spanish Type Culture Collection). BGSC (Bacillus Genetic Stock Center). DSMZ (German Collection of Microorganisms and Cell Cultures). UMA (University of Malaga).

### Genome sequencing, assembly and annotation

Bacterial DNA was isolated using the JetFlex^™^ Genomic DNA Purification Kit (Thermo Fisher Scientific, Bremen, Germany) following the kit protocol for Gram-positive bacteria. A DNA library was generated by NextSeq 550 (Illumina, United Kingdom) sequencing. The genome assembly was performed using the *B. cereus* type strain (ATCC 14579) as the reference strain of the group with the workflow of Pipeline A5-miseq [21]. The assembled genomes were annotated with the NCBI Prokaryotic Genome Annotation Pipeline (PGAP) version 4.3 algorithm [22] and deposited in NCBI GenBank (**Table S1**). Genomes of strains used in this work (ATCC 14579, AH187 and DSM 2302) were sequenced, and their genomes were obtained from the NCBI complete bacterial genome repository (http://www.ncbi.nlm.nih.gov/genome/). This whole genome shotgun project has been deposited at DDBJ/ENA/GenBank under accession number *PHKO00000000*. The version described in this paper is version *PHKO0000000*.

### Comparative genome, pangenome and phylogenetic analysis

Genomic analyses for comparison of the strains listed in **Table 1** were performed as follows: amino acid sequences for each strain were obtained from the GenBank files using the Perl programming language. Pfam domains [23] were identified inside the proteins using a Perl script and an InterProScan algorithm in the InterPro database [24]. Protein-protein comparisons among the strains of the analysis were performed to obtain the bidirectional best hits (BBH), and orthologous and paralogous proteins were identified using the NCBI BLASTP algorithm. The protein sequences were clustered according to a percentage of identity greater than 80%, which retained the same Pfam domain with the OrthoMCL algorithm [25]. The protein clusters were classified into three different groups: core (proteins shared among all the studied strains), accessory (proteins shared among all study strains except one or less) and unique (proteins only present in one of the studied strains and not in the others). To compare the function of *B. cereus* strains, GO terms (The Gene Ontology Consortium, 2015) and KO terms (KEGG Orthology) were obtained and analysed using InterProScan. The WEGO (Web Gene Ontology Annotation Plot) tool (http://wego.genomics.org.cn/) was used for GO term annotation, and the KAAS server (https://www.genome.jp/kegg/kaas/) [26] and BlastKOALA (https://www.kegg.jp/blastkoala/) [27] were used for KO term annotation. GO and KO term comparisons and visualization were performed using GraphPad Prism version 6.0 (GraphPad Software, San Diego, California, USA) and InteractiVenn [28]. Multilocus sequence phylogenetic analyses of the nine strains used for the pangenomic analysis, employing *Bacillus subtilis* subsp. *subtilis* 168 as an outgroup reference organism, was performed using the nucleotide sequences of the 16S rRNA, DNA gyrase B subunit *gyrB*, RNA polymerase sigma 70 factor *rpoD* and RNA polymerase beta 70 factor *rpoB* genes. The MUSCLE algorithm included in MEGA7 software was utilized to align the concatenated gene sequences. Next, a maximum likelihood tree was built with the MEGA7 program using the neighbour-joining substitution model, and bootstrap values of each branch were calculated 1 000 times. To study the presence/absence of sporulation genes, a comparison between the *B. cereus* AH187 and *B. cereus* DSM 2302 genomes was performed with the Gview server tool (https://server.gview.ca). The *B. cereus* AH187 gff file was obtained from NCBI (ASM2122v1), and the *B. cereus* DSM 2302 fasta file was generated in the pangenome study described elsewhere in this publication. Unique genes were used to identify the Gene Ontology functional categories using comparativeGo (https://www.comparativego.com). Sporulation genes from these two strains were selected and compared by blast2 sequencing to obtain the final list of sporulation genes of *B. cereus* AH187 with no matches in *B. cereus* DSM 2302. To determine the possible function of genes annotated as hypothetical proteins or with unknown function, their amino acid sequences were compared with proteins from other organism using the HHpred tool of the Max Planck Institute for Developmental Biology (Tübingen, Germany) [29, 30]. We also searched for orthologous genes in other *Bacillus* species, or we determined the hypothetical function according to the corresponding GO/KO term of the Pfam domain.

### Sporulation and image acquisition

The sporulation assays were performed in DSM medium. One colony of each strain was picked from an LB agar plate and resuspended in DSM broth. Each strain was incubated at 28°C with shaking for 24 h. Bacterial concentrations in CFU/ml were determined by plating 100 µl aliquots of 10-fold serial dilutions onto LB agar. To estimate the proportion of spores, each dilution was heated at 80°C for 10 min and subsequently plated. Microscopy images from DSM cultures were taken on a Nikon Eclipse Ti fluorescence microscope with a Super Plan Fluor objective 100x equipped with a Hamamatsu Orca R2 monochrome 1.3 MP CCD camera. Image processing (for proportion of spores) was performed using ImageJ software.

### Construction of a *B. cereus* sporulation mutant

A *B. cereus sigE* mutant was obtained by electroporation using the pMAD plasmid [31]. To generate mutagenesis constructs, NEBuilder HiFi DNA Assembly Master Mix (New England Biolabs) was utilized following the manufacturer’s instructions, and specific primers (**Table S2**) were employed to amplify the upstream and downstream regions of the *sigE* gene. The pMAD vector was digested with SmaI and incubated at 50°C for 1 h in a thermocycler with the upstream and downstream fragments. The total amount of fragments was 0.2 pmol with a 1:2 DNA ratio (vector:insert). The pMAD-*sigE* plasmid was utilized to transform electrocompetent *B. cereus* cells by electroporation as described previously with several modifications [17, 32]. One microgram of plasmid was incubated with 100 µl of electrocompetent cell suspension on ice for 5-10 min. The mixture of bacteria and DNA was electroporated in a 0.2-cm chilled cuvette using the following electroporation parameters: 1.4 kV voltage, 25 µF capacitance and 200 Ω resistance. The electroporated cells were recovered in LB at 30°C for 5 h and then plated onto LB plates supplemented with erythromycin to select bacterial transformants. To obtain mutants, *B. cereus* transformants were incubated at 40°C without antibiotics to block plasmid replication and force the shuttle vector to integrate into the chromosome via homologous recombination.

### Dynamics of bacterial cells on plants

To evaluate the persistence of *B. cereus* strains on plants, melon (*Cucumis melo* cv. Rochet Panal) (Fitó, Barcelona, Spain) leaves, cucumber (*Cucumis sativus* cv. Marketer) (Fitó, Barcelona, Spain) leaves, cucumber fruit and endive (*Cichorium endivia*) were analysed. Melon and cucumber plants were grown under greenhouse conditions, and fresh cucumber fruit and endive were purchased in a grocery store. Cucumber fruit and endive were rinsed with 1% (w/v) sodium hypochlorite and 96% ethanol in distilled water for 4 min before being inoculated with *B. cereus* cell suspensions. A bacterial suspension (1 ml) at a concentration of 10^8^ CFU/ml in distilled water was spread onto the adaxial axis of second and third leaves of melon or cucumber leaves and around cucumber fruit and endive leaves. After bacterial inoculation, the melon and cucumber plants were incubated in a growth chamber at 25°C with a 16-h light/8-h dark photoperiod and 85% relative humidity. Endive and cucumber fruit were incubated at room temperature. Bacterial persistence and sporulation were evaluated 15 days after inoculation. Leaves and cucumber fruit skin were placed individually into sterile plastic stomacher bags with 10 ml of distilled water and homogenized for 3 min in a stomacher homogenizer (Interscience, France). One hundred microlitres of 10-fold serial dilutions of the homogenates were plated onto LB agar plates and incubated at 28°C for 24 h. To estimate the level of spores, serial dilutions were incubated at 80°C for 10 min to kill vegetative cells, and the spores were subsequently plated and incubated overnight at 28°C.

### Biofilm and swarming motility assays

*In vitro* biofilm assays of *B. cereus* strains were performed in TY broth and in TY 1.5% agar supplemented with 20 µg/ml and 10 µg/ml of Congo Red and Coomassie Brilliant Blue filtered dyes, respectively [17, 33] For biofilm assays in TY broth, a colony from a pre-culture in LB agar at 28°C for each strain was previously resuspended in 1 ml of TY medium; 10 µl of this culture was inoculated into 1 ml of TY broth in the center wells of a 24-well plate, and the plate was incubated without shaking at 28°C for 72 h. To evaluate bacterial adhesion to abiotic surfaces (i.e. adhesion to the well wall), the biofilm of each well was stained with 1% crystal violet solution [34] for 3 min and subsequently rinsed several times with distilled water until the excess dye was removed. Finally, the plate was dried on filter paper on a bench. To study biofilm formation on agar plates (colony morphology and staining), 2 µl of the preculture described above was inoculated separately on 1.5% TY agar plates with Congo Red and Coomassie Brilliant Blue, and the plates were incubated at 28°C for 72 h. To study swarming motility, 0.7% TY agar plates were utilized. Two microlitres of the pre-culture described above was inoculated on TY plates and incubated at 28°C for 72 h before measuring the diameter of colonies.

### Leaf and fruit metabolome analysis

Metabolomic analysis of melon and endive leaves was performed by Metabolon^®^ (Durham, North Carolina, USA). One hundred milligrams of each tissue (from a pool of leaves) was homogenized and lyophilized. Samples were prepared using the automated MicroLab STAR^®^ system from Hamilton Company (Franklin, Massachusetts, USA). Samples were extracted with methanol under vigorous shaking for 2 min to precipitate protein and dissociate small molecules bound to protein or trapped in the precipitated protein matrix followed by centrifugation to recover chemically diverse metabolites. The resulting extract was divided into four fractions: two for analysis by two separate reversed-phase (RP)/UPLC-MS/MS methods using positive ion mode electrospray ionization (ESI), one for analysis by RP/UPLC-MS/MS using negative ion mode ESI, and one for analysis by HILIC/UPLC-MS/MS using negative ion mode ESI. Samples were placed briefly on a TurboVap® (Zymark) to remove the organic solvent. The metabolic analysis was performed by ultra-performance liquid chromatography (UPLC) coupled to a mass spectrometer with a heated electrospray ionization (HESI-II) source and Orbitrap mass analyser operated at 35,000 mass resolution. The four extracts were dried and subsequently reconstituted in methanol. One aliquot was analysed using acidic positive ion conditions, chromatographically optimized for more hydrophilic compounds and subsequently gradient-eluted from a C18 column (Waters UPLC BEH C18-2.1×100 mm, 1.7 µm) using water and methanol containing 0.05% perfluoropentanoic acid (PFPA) and 0.1% formic acid (FA). A second aliquot was also analysed using acidic positive ion conditions but was chromatographically optimized for more hydrophobic compounds. The extract was gradient-eluted from the aforementioned C18 column using methanol, acetonitrile, water, 0.05% PFPA and 0.01% FA and was operated at an organic content that was higher overall. A third aliquot was analysed using basic negative ion optimized conditions using a separate dedicated C18 column. The basic extracts were gradient-eluted from the column using methanol and water but with 6.5 mM ammonium bicarbonate at pH 8. The fourth aliquot was analysed via negative ionization following elution from a HILIC column (Waters UPLC BEH Amide 2.1×150 mm, 1.7 µm) using a gradient consisting of water and acetonitrile with 10 mM ammonium formate, pH 10.8. The MS analysis alternated between MS and data-dependent MS^n^ scans using dynamic exclusion. The scan range varied slightly between methods but covered approximately 70-1000 m/z. Raw data were extracted, peaks were identified, and data were submitted to QC processing using Metabolon’s hardware and software.

To visualize the differences between melon and endive leaves, MetaboAnalyst (https://www.metaboanalyst.ca) was used to generate the heatmap with hierarchical clustering and to determine the most important metabolites through PLS-DA analysis.

### Construction of artificial leaf surfaces, bacterial inoculation and incubation

To produce artificial leaf surfaces, melon leaves were grown as described above. The second and third leaves were taken to mould the leaf surface replica. The process of making artificial leaves was undertaken as previously described [35] and summarized as follows: first, a negative stamp of the adaxial part of the leaf was created. The negative stamp was made with a mixture of 10:1 polydimethylsiloxane (PDMS) and curing agent from a Sylgard 184 PDMS elastomer kit (Dow Chemical, Midland, USA). The mixture was poured over the real leaf previously joined with double-sided adhesive tape on a Petri dish and incubated at 30°C for 17-24 h. When the negative stamp was solidified, it was carefully separated from the real leaf and subsequently exposed to UV light (400 F, 400 W and λ = 187-254 nm) for 1 h to fix the PDMS. After UV exposure, the negative stamp was treated with a mixture of 1000:1 toluene and octadecyltrichlorosilane (ODTS) for 5 min to create a soft cover that facilitated the later separation of the negative stamp from the positive stamp. Next, the negative stamp was rinsed with absolute ethanol for 2 min and dried for at least 10 h at room temperature. To make the positive stamp (artificial leaf), the PDMS mixture was made as described above, and it was poured onto the negative stamp, which was attached to a Petri dish. When the PDMS mixture was solidified, the positive stamp was carefully separated from the negative stamp. The fabricated artificial leaves were autoclaved and rinsed with a mixture of 1:1000 Triton X-100 in distilled water. Artificial leaves were cut into coupons measuring 25.75 cm in diameter and inoculated with 20-µl drops (5 drops per coupon) of a 10^8^ CFU/ml bacterial suspension in distilled water [35, 36] or in distilled water supplemented with 0.5% quinate, 0.1% galactarate and 0.1% saccharate or with 1 mM isovitexin to mimic endive and melon leaf metabolite profiles, respectively (obtained from the metabolomic analysis). Inoculated coupons were incubated over moistened filter paper with 10 ml of distilled water into a Petri dish plate or without humidity conditions (without distilled water), allowing the drops to dry on an open Petri dish. All the plates with coupons were incubated for 15 days in a growth chamber as described above for real leaves. After 15 days, the coupons were vortexed for 15 sec and sonicated for 10 min in an ultrasonic bath in 10 ml of distilled water in a 50-ml tube [37]. Tenfold serial dilutions were made, and 100 µl of each dilution was plated on LB agar. To estimate the level of spores, the dilutions were heated at 80°C for 10 min and plated and incubated at 28°C for 24 h to obtain CFU counts.

### Gastric passage simulation

To analyse gastric passage simulation after food ingestion, we employed the protocol described previously [20, 38]: *B. cereus* strains were incubated in LB agar at 30°C for 24 h. One colony was transferred into a 250-ml flask with 40 ml of J broth (JB) and incubated for 18 h at 30°C and 130 rpm. *B. cereus* cells were added to a 1:1 mixture of JB and gastric media (GM) (4.8 g/l NaCl, 1.56 g/l NaHCO_3_, 2.2 g/l KCl and 0.22 g/l CaCl_2_) at pH 2 and 4.5 plus 500 U/l pepsin to an initial concentration of 10^6^-10^7^ CFU/ml bacteria and incubated at 37°C for 5 h and 130 rpm to simulate human stomach conditions. Samples were taken hourly, 10-fold serial dilutions were made in 0.2 M sodium phosphate buffer at pH 7 to neutralize the stomach pH, and 100 µl of each dilution were plated onto LB agar. For the determination of spores, the dilutions were heated at 80°C for 10 min and plated onto LB agar.

### Inoculation of mice with *B. cereus* and histological analysis of intestines

CD1 female mice (3 months old) were purchased from the Andalusian Centre for Nanomedicine and Biotechnology (University of Malaga). Each experimental group consisted of 6 mice housed in cages that were maintained at a controlled temperature and light cycle; the mice were fed an autoclaved standard balanced conventional diet and a gelatine-only diet 24 h before the infection. Animals were handled in accordance with international regulations for animal welfare. Bacteria were administered by intragastric gavage using a blunt-end needle. Six randomly allocated mice were challenged with 100 ml of 10^8^ CFU of *B. cereus* DSM 2302 in 1x PBS or with 100 µl of 1x PBS [39]. After 6 h of bacterial administration, mice were anaesthetized with an injection of 200 ml of a mix of 10% ketamine, 6% xylazine and 84% water each and sacrificed by cervical dislocation. Samples from the caecum region were aseptically collected and washed in PBS and immediately fixed in 4% formalin overnight (4°C) for histological preparation. To obtain frozen sections, tissues were cryoprotected in 15% and 30% sucrose:PBS solutions, embedded in OCT, snap-frozen in liquid N_2_-cooled isopentane and sectioned in a cryostat. OCT blocks were cut into 10-mm sections, and some of them were stained with haematoxylin–eosin. At least three caeca were analysed for each treatment.

### Immunohistochemistry and confocal microscopy

Immunofluorescence analyses were performed by blocking nonspecific binding sites with 16% sheep serum, 1% bovine serum albumin and 0.5% Triton X-100 in PBS (SBT) and incubating either tissue section slide with primary antibodies (anti-cytokeratin “DAKO Z0622, 1:100 dilution” and anti-PanCadherin “Sigma C3678, 1:100 dilution”) overnight at 4°C. Next, the samples were washed in PBS (3 × 5 min) and incubated with the secondary antibody (Alexa Fluor® 647 “InmunoJackson 711-605-152”, 1:200 dilution) + DAPI (Sigma D9542, 1:2000 dilution) for 2 h at room temperature. All samples were finally washed in PBS (3 × 5 min), mounted in a 1:1 PBS/glycerol solution and analysed using an SP5 laser confocal microscope (LEICA).

### Ethics approval and informed consent for mouse experiments

Experiments were carried out in accordance with the Spanish Institutional Animal Use and Care Committee regulations and with the approval of the Regional Andalusian Government (code A/ES/14/43). The methods were carried out in accordance with the relevant guidelines and regulations.

### Statistical analysis

The statistical analyses were performed using the statistical software GraphPad Prism version 6.0 (GraphPad Software, San Diego, California, USA). For melon and endive leaf experiments, one-way nonparametric ANOVA (Kruskal-Wallis test) was performed with Dunn’s multiple comparison tests. For artificial surface experiments, two-way ANOVA (with the Bonferroni test) was applied. The results with p-values < 0.05 were considered to be significant. Metabolome data were normalized for internal consistency by processing a constant weight of sample per volume of extraction solvent. No *post hoc* mathematical normalization was imposed on the data. Data were scaled to the median value for each compound, and missing values were imputed with the minimum detected value for that compound. Statistical calculations were performed using natural log-transformed scaled imputed data (table not included).

## Results

### Sporulation permits *B. cereus* to persist on melon leaves but is dispensable on endive

We investigated the ability of a group of *B. cereus* strains isolated from different outbreaks of food poisoning (**Table 1)** to colonize and persist on leave of two different plant species, melon and endive. Two differentiated behaviours of *B. cereus* strains were observed on and in melon leaves: seven strains (AH187, ATCC 14579, DSM 4282, DSM 4313, DSM 14729, UMAF8564 and 6A27) were capable of persisting at 10^4^-10^6^ CFUs per gram of leaf and had a spore content of 20-40% (**Fig. S1**), and two strains (DSM 8438 and DSM 2302) had CFUs per gram and spores counts below the detection threshold (**Fig. S1**). In endive, however, all the strains persisted at 10^4^-10^6^ CFU per gram of leaf. The percentages of sporulation were similar to those recorded on melon leaves with the exception of DSM 2302, which did not show any sign of sporulation. With these findings, we compared the population dynamics of strain AH187 (a representative of the group of persister and sporulating strains), DSM 8438 and DSM 2302 (nonpersister on melon leaf and nonsporulating strains) on both plant leaves (**Fig. 1**). The population size of AH187 progressively decreased and stabilized at 10^6^ CFU per gram of melon leaf 8 days after inoculation; however, DSM 8438 and DSM 2302 showed a sharp decrease within the first 4 days and became undetectable after 8 days of inoculation (**Fig. 1)**. The AH187 population dynamics on endive leaves were similar to those on melon leaves, and DSM 8438 and DSM 2302 stabilized at 10^4^-10^5^ CFU per gram of endive leaves after 2 DAI (**Fig. 1)**. The sporulation rate of AH187 and DSM 8438 increased starting at 4-6 HAI, and no spores were detected for DSM 2302. Similar results for the persister group were recorded on cucumber leaves and fruit, and DSM 8438 and DSM 2302 did not sporulate in any of the studied cucumber organs (**Fig. S1**).

**Figure 1.**
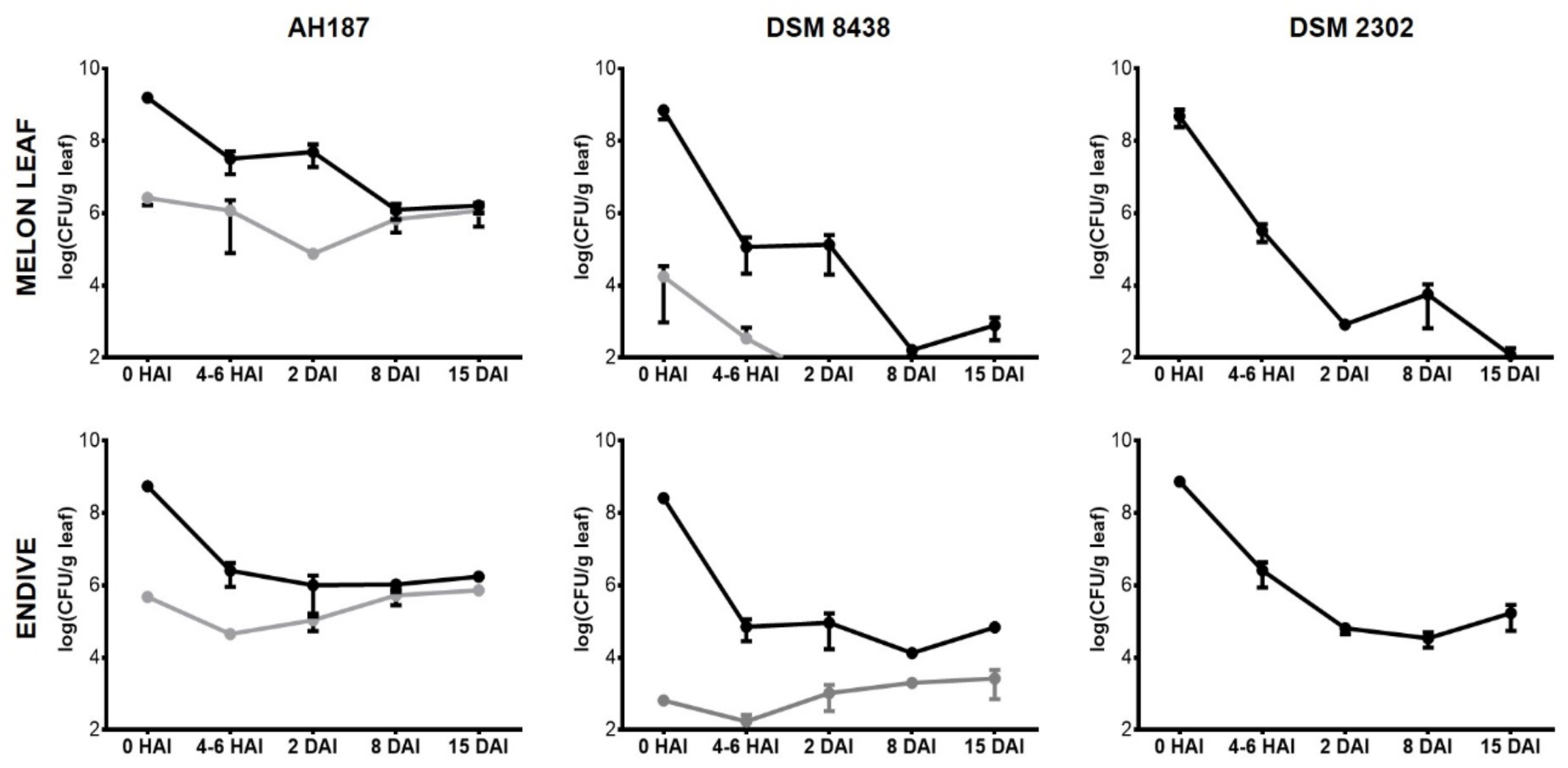
Sporulation is essential for the persistence of *B. cereus* on melon leaves but not on endive leaves. CFU counts per gram of melon and endive leaf after inoculation (0 hours (HAI) 4-6 h (HAI), 2 days, 8 days and 15 days (DAI)) with *B. cereus* strains AH187, DSM 8438 or DSM 2302 strains. The top plots correspond to melon leaf counts, and the bottom plots correspond to endive counts. The black line shows the counts of total CFUs on melon and endive leaves. The grey line shows the CFU counts corresponding to spores on melon and endive leaves. The detection limit of the technique is 10^2^ CFU/g leaf. Error bars are the standard error of three or more replicates.

### *B. cereus* DSM 2302 is genetically impaired in sporulation but capable of biofilm formation

We sequenced and compared the genomes of all the strains used in this study to define the genetic defects for the inability of DSM 2302 to sporulate, and we searched for genetic traits defining a plant-associated lifestyle. Multilocus sequence analysis using 4 conserved genes (16S rRNA, *gyrB, rpoB* and *rpoD*) demonstrated that AH187, DSM 2302 and DSM 8438 belonged to completely separate phylogenetic branches (**Fig. S2**). This finding enabled us to surmise that the strains that do not share specific niches employ different evolutionary strategies and that their evolutionary distance might be related to their different lifestyles. A deeper study of the pangenome of the nine *B. cereus* strains (**Fig. S3A**) identified a total of 48 844 coding DNA sequences (CDSs) in the nine *B. cereus* strains. Of these total CDSs, 30 746 were highly conserved across all nine genomes and comprised the core genome of these *B. cereus* strains. The core genes were clustered into 3 326 orthologous groups (OGs), the additional genes (see material and methods for details) were clustered into 3 325 Ogs, and the unique genes of the 9 strains comprised 3 050; therefore, all CDSs were clustered into 9 702 Ogs. Notably, AH187 and DSM 2302 had the largest number of unique genes, exhibiting a total of 444 and 414 exclusive sequences, respectively, which supported our hypothesis on the specialization of these strains to different ecological niches (**Table S3**). Further analysis of the core genome by 10 different combinations of the sequences of the nine studied strains demonstrated that the number of genes did not decrease with the increase in the number of genomes added to the study (**Fig. S3B**). In addition, the pangenome of the studied strains was open, indicating that each sequenced genome contributed new genes to the pangenome but never reached an exact number of genes (**Fig. S3B)**. A more specific comparison of DSM2302 with AH187 showed an uneven distribution of the genes or the absence of several complete regions, including operons (**Fig. S3C)**, as well as widely conserved regions in both strains.

Consistent with the apparent phenotypic defect in sporulation exhibited by DSM 2302, 36 genes involved in the sporulation programme were absent (**Table 2, Fig. 2D**) [40, 41, 42]. To confirm this genotype, we analysed the intrinsic ability of the strains to sporulate in the spore-promoting medium DSM (**Fig. 2B**). All the strains that could persist over melon leaves showed high sporulation rates in this medium (**Fig. 2B, grey bars**), but no sign of sporulation was observed in cultures of DSM 2302. Contrast-phase microscopy analysis of single cells confirmed the presence of refringent endospores within AH187 cells but neither endospores nor mother cell-released spores in cultures of DSM 2302 (**Fig. 2B)**.

**Table 2.**
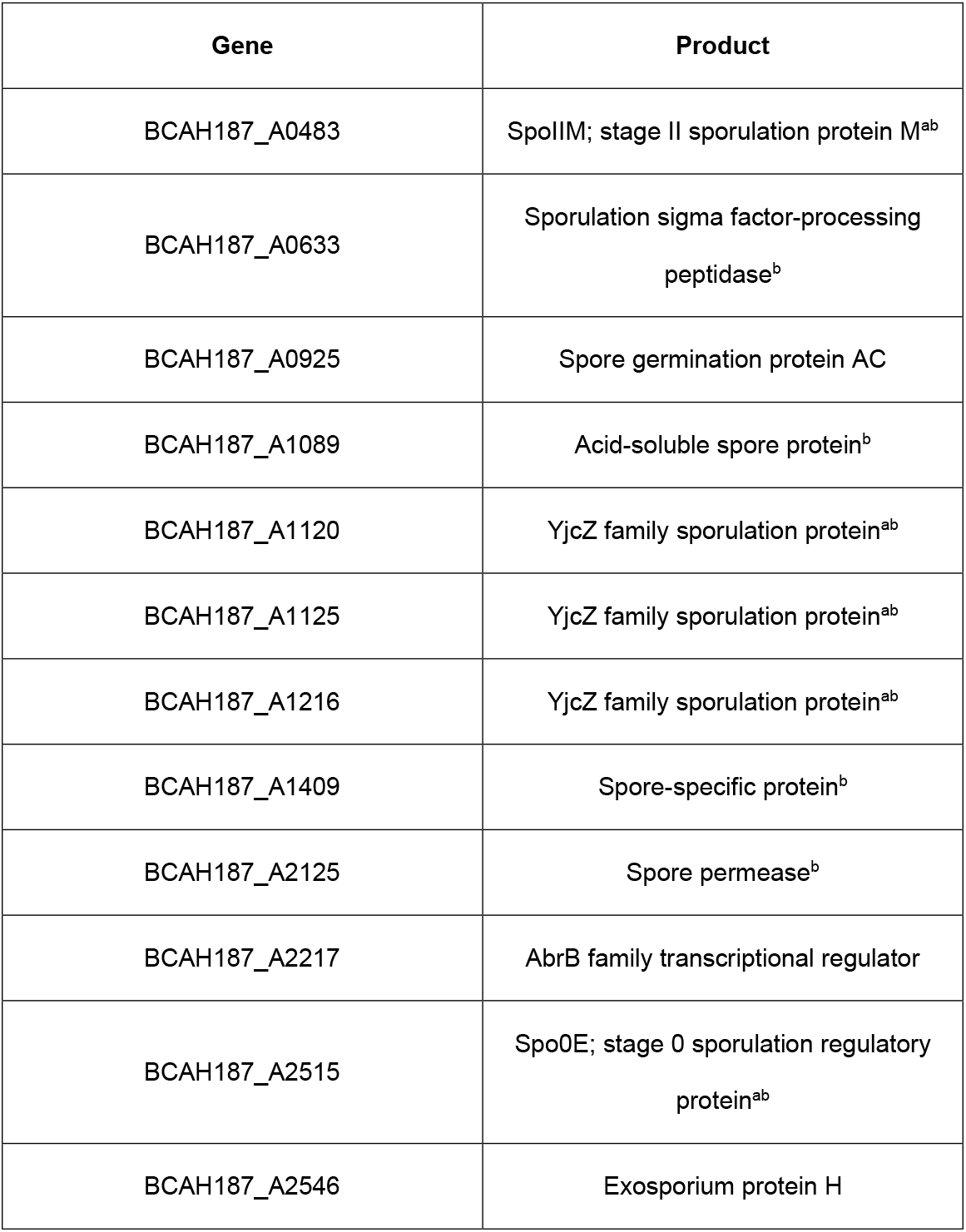
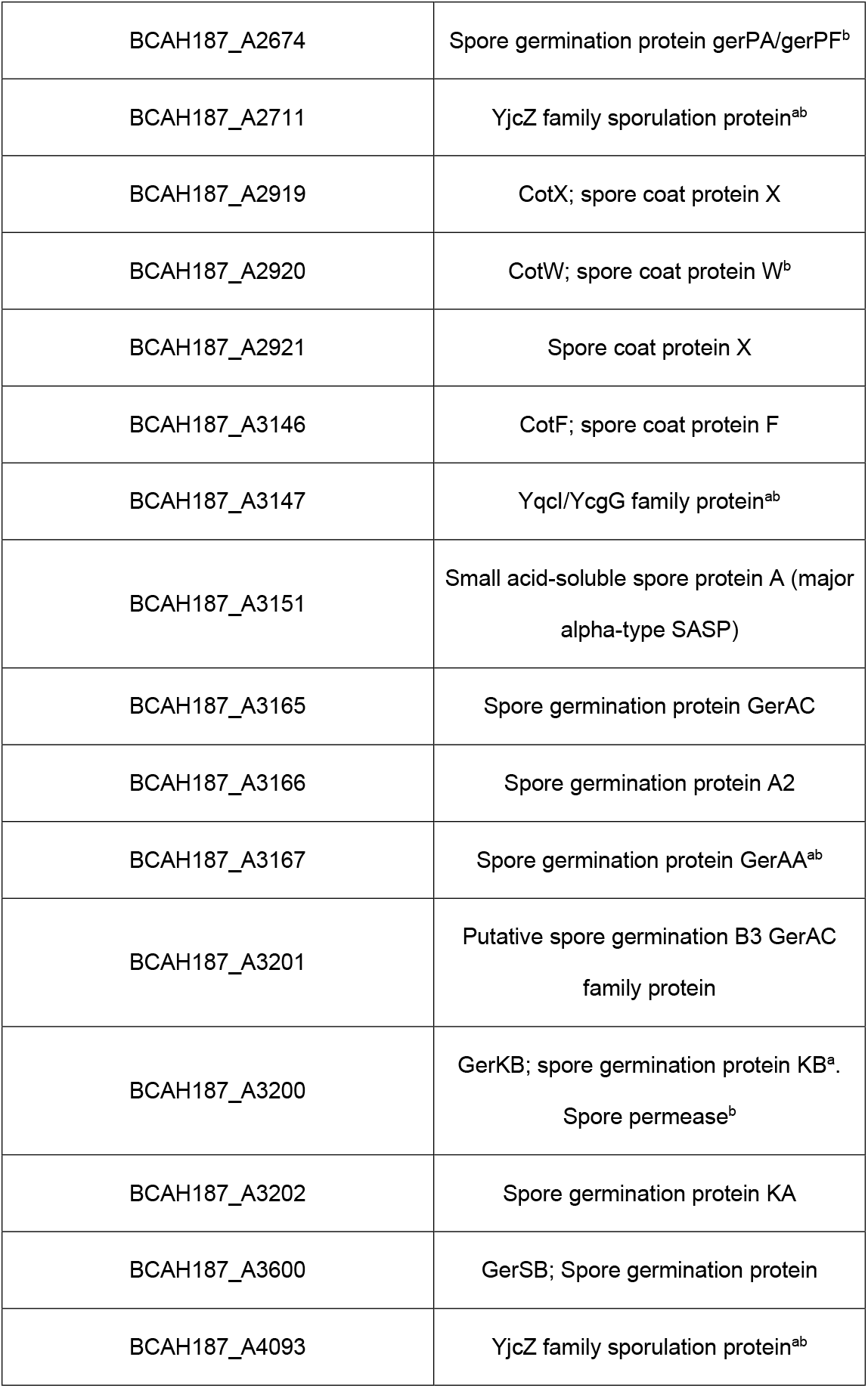
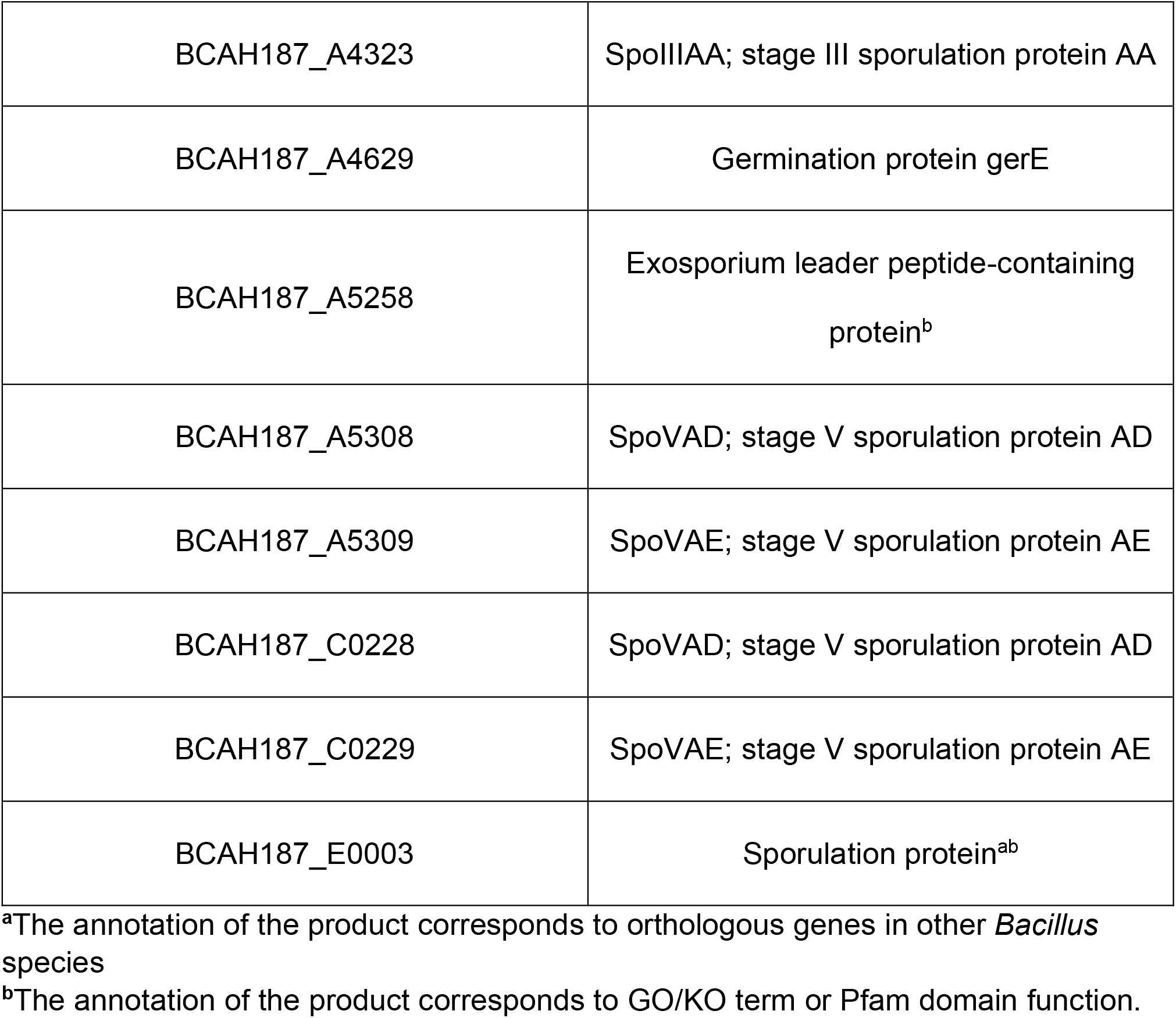
Genes absent in DSM 2302 strain compared with AH187 strain. Gene corresponds to AH187 locus absent in DSM 2302 strain. Product indicates the protein associated to each gene.

**Figure 2.**
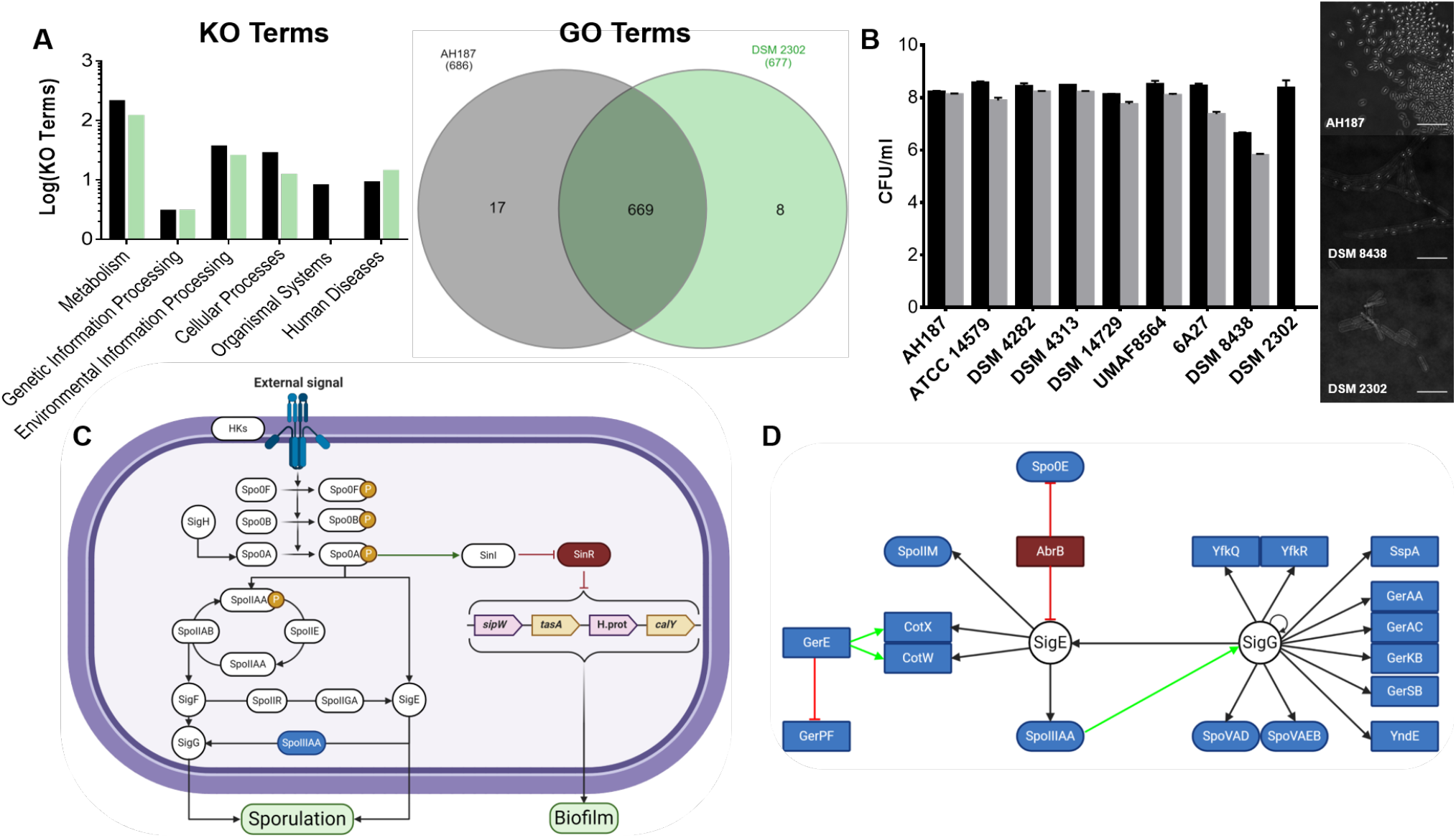
*B. cereus* DSM 2302 is unable to sporulate, but it retains the machinery to form biofilms and cause human diseases. **A** KO and GO term comparison between AH187 and DSM 2302 strains. Black bars and grey circles correspond to the AH187 strain, and green bars and circles are the DSM 2302 strain. The KO plot shows unique KO terms of each strain compared to the other. Pie plot showing GO terms shared and nonshared between AH187 and DSM 2302 strains. See also Table S3, which shows KO and GO terms present in DSM 2302 and not in the AH187 strain. **B** Sporulation of *B. cereus* strains in DSM medium after 12 h incubation at 30°C with shaking. The plot shows the counts of total CFU/ml (black bars) and spores (grey bars) plated in LB agar after incubation in DSM medium. The detection limit of the technique is 10^1^ CFU/ml. Error bars are the standard deviation of three or more replicates. Statistical analysis is the result of the comparison of all strains against AH187. * (P<0,05), ** (P<0,01), *** (P<0,001). The pictures correspond to microscopy images showing the different sporulation behaviours of the AH187, DSM 8438, and DSM 2302 strains after 12 h of incubation at 30°C with shaking in DSM medium. Scale bar corresponds to 10 µm. **C** Sporulation and biofilm pathway in *B. cereus* group. Blue proteins are absent in DSM 2302 strain. **D** Sporulation pathway of genes absent in *B. cereus* DSM 2302. Blue proteins refer to genes present in AH187 and absent in strain DSM 2302 in addition to AbrB regulator. Green arrows indicate an activation process, and red arrows indicate repression of the target gene. SigE and SigG are present in DSM 2302 genome.

In *Bacillus*, sporulation and biofilms are known to contribute to the persistence of *Bacillus* and other species in different habitats, and these two cellular destinies are under the control of the master regulator *spo0A* (**Fig. 2C**). We disregarded a hypothetical collateral interruption of the biofilm regulatory pathway in DSM2302, given that most of the lacking genes related to sporulation were structural and belonged to stages of the sporulation programme downstream of phosphorelay (**Table 2**). In addition, we identified the two major regulators of the expression of biofilm-related genes *sinR* and *sinI* along with the gene cluster *sipW*-to-*calY*, known to be essential for the assembly of the extracellular matrix (ECM) (**Fig. S4B**). Another indispensable component of the ECM is exopolysaccharides (EPSs). Two genomic regions containing EPS-related genes were found in DSM 2302: i) one similar to the so-called *eps1* of the type strain ATCC 14579 and orthologous to the *epsA*-*O* of *B. subtilis*. This region was also found in strain AH187 which, notably, varied in the genes of the central core of the region (**Fig. S4C**). ii) One region homologous to the so-called *eps2* region of ATCC 14579, which was absent in strain AH187 (**Fig. S4D**). According to this genetic information, DSM 2302 was able to form biofilms in static cultures, although phenotypically differentiated from AH187. Colonies of DSM 2302 stained with Congo Red (known to be related to the product of the *eps2* region in the type strain ATCC 14579) and adhered to the walls and bottom of the wells but failed to form pellicles in the air-liquid interphase of static liquid cultures. Strain AH187 formed colonies that stained slightly with Congo Red, formed pellicles in the air-liquid interphase and adhered to the wells in static liquid cultures. (**Fig. S4A**). The pangenomic analysis of DSM 2302 additionally showed several genes involved in the assembly of the ECM, polysaccharide and putrescine biosynthesis, all of which are putative structural integrant of the biofilm programme (**Table S4)**.

Further genomic analysis demonstrated the presence of genes related to a plant-associated lifestyle in the genome of AH187 but absent in DSM 2302. These genes included several response regulators; ABC transporters hypothetically involved in exoprotein production, competence or sporulation; a regulator inducible by reactive oxygen species (ROS); a putative endoglucanase involved in plant cell wall degradation; diverse surface proteins and important enzymes related to plant nitrogen sources, such as asparagine, glutamine or proline (**Table S5**). In contrast, DSM 2302 had more genes related to human disorders, bacterial invasion of epithelial cells, bacterial extracellular infection, cell wall surface anchoring, cell adhesion and internalization, and antimicrobial resistance in the genome (**Fig. 2A**), most of which were observed in the related food-borne bacteria *Listeria monocytogenes* and *Escherichia coli* or *Yersinia enterocolitica* [43, 44].

In summary, the defect of sporulation, the lack of genes related to the metabolism of nitrogen sources related to plants and the overrepresentation of genes associated with pathogenic interactions with humans are consistent with the low survival of DSM 2302 on plants and with an ecological adaptation to human-related niches.

### Metabolic nature of endive enables the persistence of vegetative cells of *B. cereus*

In addition to the lack of sporulation, the different fitness levels of DSM 2302 on melon or endives leaves raised the question of the major influence of the host. With in-depth metabolomic analysis of leaf tissue, the chemical profiles of both plant species demonstrated significant differences in metabolic families and metabolite concentrations (**Fig. 3A**). Carbohydrates, amino acids, lipids and nucleotides were largely observed in the endive (**Fig. 3B**). More specifically, chlorogenate, allantoic acid, quinate, galactarate and glucarate were clearly the most representative endive metabolites. However, secondary metabolites of flavonoids (such as isovitexin), glutathione, pinoresinol and siderophores were the most frequently observed metabolites in melon leaves (**Fig. 4C**).

**Figure 3.**
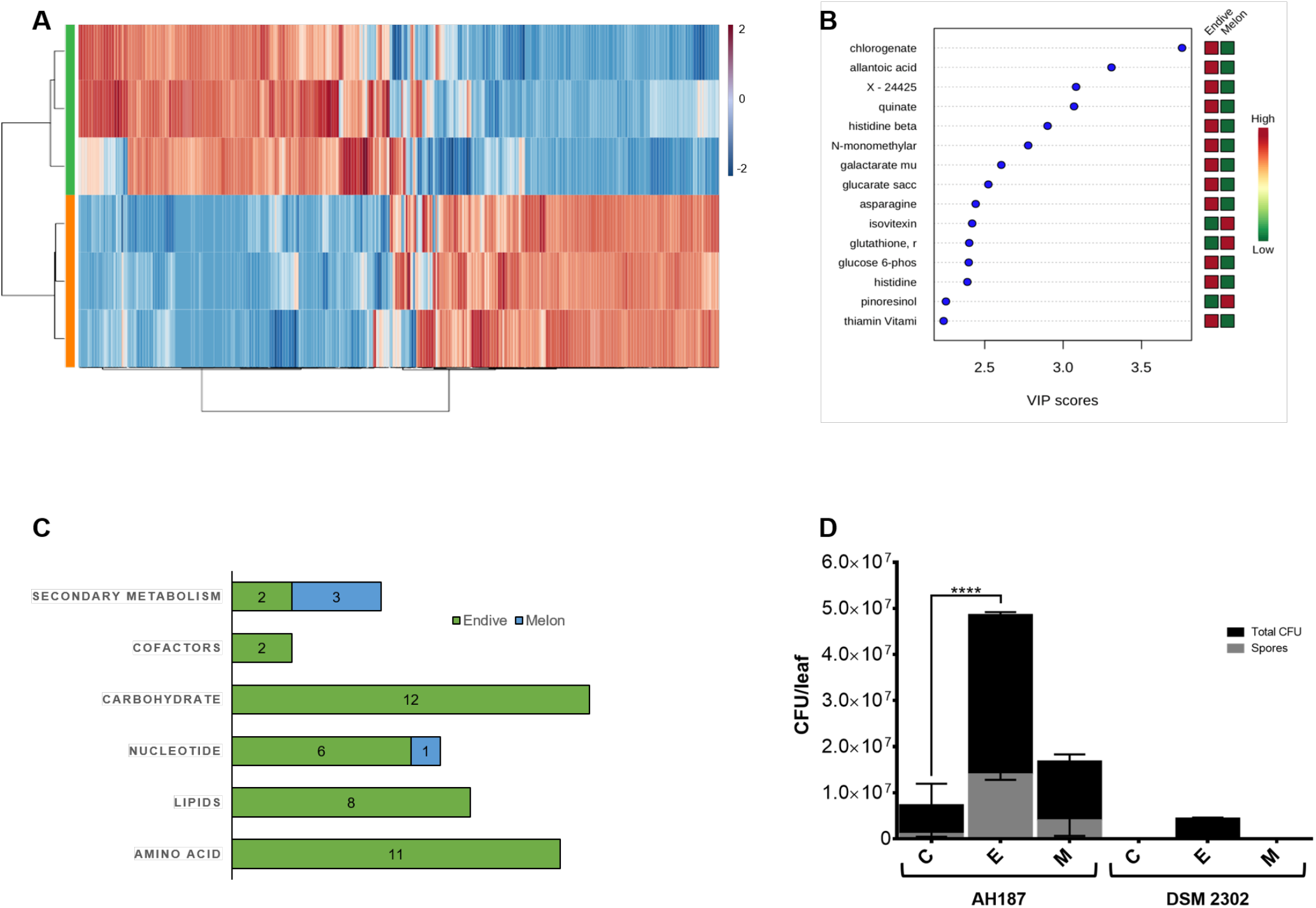
Differences in the nutrient content of the plant host impact the survival of non-sporulating *B. cereus* DSM 2302. **A** Clustering result shown as a heatmap of three replicates of leaves of endive (orange) and melon leaves (green). In the heatmap, red indicates a higher concentration of a metabolite; in contrast, blue indicates a lower concentration of the metabolite. **B** Major metabolites grouped by superfamilies found in melon and endive leaves. **C** Important metabolites identified by PLS-DA. The coloured boxes on the right indicate the relative concentration of the corresponding metabolite in each studied surface (red indicates a higher relative concentration, and green indicates a lower relative concentration of the corresponding metabolite). **D** CFU counts per artificial leaf surface: without metabolites (C) with the majority endive metabolites (quinate, galactarate and glucarate) (E) and the majority flavonoid (isovitexin) found in melon leaves (M) 15 days after inoculation with *B. cereus* AH187 and DSM 2302 strains. The black bar corresponds to the total CFUs per artificial leaf surface, and the grey bars are the counts of CFUs corresponding to spores. The detection limit of the technique is 10^2^ CFU/g leaf. Error bars are the standard error of three replicates. Statistical analysis is the result of the comparison of the DSM 2302 strain against AH187. * (P<0.05), ** (P<0.01), *** (P<0.001), ****(P<0.0001).

**Figure 4.**
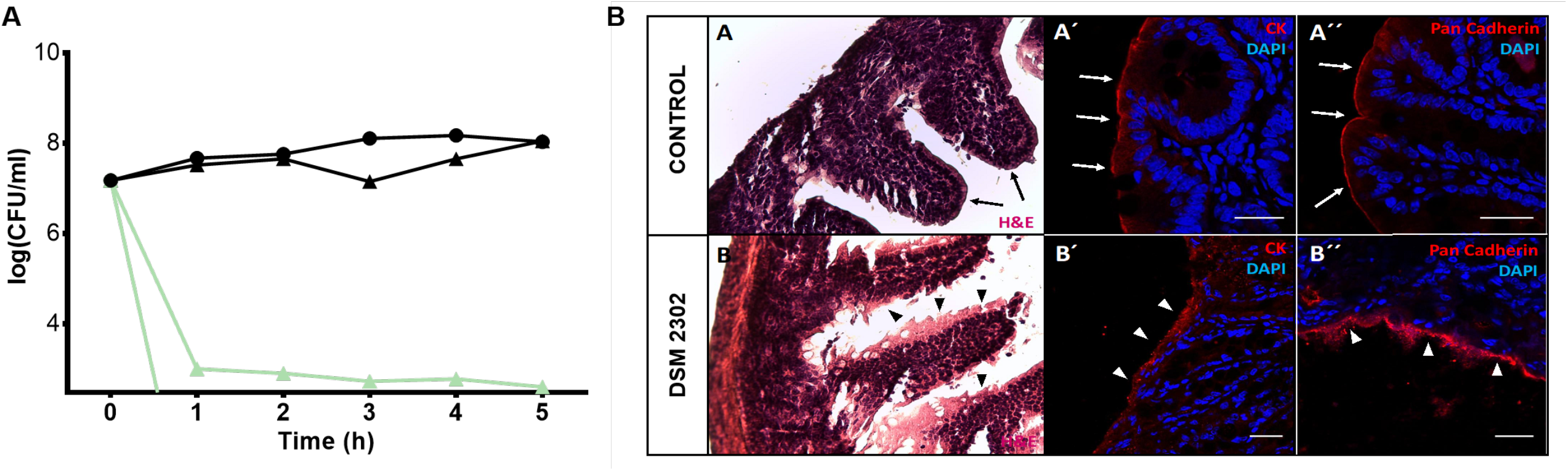
Vegetative cells of DSM 2302 can survive simulating stomach conditions and cause symptoms in mouse intestines. **A** Counts of total CFU/ml on gastric media (GM) supplemented with JB broth of AH187 (▲) and DSM 2302 (●) strains at 37°C for 5 h. Black lines correspond to pH 4.5 simulating conditions after food intake. Green lines correspond to pH 2 (basal stomach conditions). The detection limit of the technique is 10^1^ CFU/ml. Error bars are the standard error of three replicates. **B** Comparative histological caecum analysis of intestine sections from control and infected mice. Haematoxylin–eosin staining showed (A) epithelium continuity in the villi and crypts of the intestinal microvilli in control intestine (PBS) and (B) epithelium disruption of intestines of infected mice. Immunohistochemistry and confocal microscopy showed epithelial continuity in the villi and crypts of the intestinal microvilli with CK (A′; arrows) and pan-cadherin (A′′, arrows) markers of the control intestine and epithelial disruption in the infected caecums using CK (B′; arrowheads) or pan-cadherin (B′′, arrowheads). All nuclei were stained with DAPI for immunofluorescence. Scale Bars (A′, A′′, B′, B′′) = 25 mm.

These findings along with the bacterial population dynamics showed above suggested that endive leaves were more conducive to bacterial growth, while melon leaves represented a more aggressive and stressful environment for bacteria. To confirm this hypothesis, we studied the bacterial behaviour in PDMS artificial surfaces that mirrored the topology of melon leaves but were supplemented with a solution of quinate, galactarate and saccharate (5:1:1), the major components detected in endive. AH187 persisted on melon artificial leaves alone or supplemented with endive metabolites; however, DSM 2302 only persisted in the presence of endive-related metabolites (**Fig. 3D**). The addition of isovitexin, the main flavonoid detected in melon leaves, permitted a slight, though insignificant, increase in the AH187 population but did not support any growth of DSM 2302 (**Fig. 3D**). The spore content of AH187 remained unchanged at 30-40% under experimental conditions (**Fig. 3D**). According to these findings, a null *sigE* (a mother cell-specific transcription factor of early sporulation in *Bacillus* spp.) mutant of AH187, unable to sporulate but capable of forming biofilms, could not sporulate or persist on melon leaves or on artificial leaf surfaces lacking the major endive-related metabolites.

### Vegetative cells of DSM 2302 can reach the intestine in mice and cause illness

The strain DSM 2302 was isolated in a food poisoning outbreak, and we have demonstrated that vegetative cells can persist on endive, a produce that is directly consumed by humans. This evidence suggests that vegetative cells of this strain may reach humans and provoke food poisoning. To test this idea, we simulated a stomach passage with basal pH conditions (pH 2) and after meal ingestion conditions (pH 4.5) using a nutritive broth (JB) as a food source. Only AH187 survived at pH 2, and this strain was observed at very low levels and largely existing as spores (**Fig. 4A**). The population densities of AH187 or DSM 2302 were comparable and remained unchanged at a pH value of 4.5 for 5 h. Thus, vegetative cells of DSM 2302 should be able to overcome the acidic condition of the stomach after food ingestion and reach the intestine. To confirm this possibility, we inoculated 10^8^ CFU of DSM 2302 into 3-month-old mice. One-third of mice inoculated with DSM 2302 cells died after 6 h of bacterial inoculation. We evaluated the histological damage of the intestine of the mice as indirect evidence of the presence of this strain in the guts. Haematoxylin–eosin staining showed continuity of the epithelium in the villi and crypts of the intestinal microvilli in untreated mice (**Fig. 4B**). However, the mice inoculated with DSM 2302 showed disruption of the epithelium. Similar results were obtained in immunohistochemical analyses using anti-Ck or anti-pan-cadherin antibodies and confocal microscopy (**Fig. 4B**).

These results enable us to conclude that vegetative cells of DSM 2302 (which lack of the acid resistance cereulide toxin) are capable of passing through the stomach and reaching the gut, subsequently causing illness in the host.

## Discussion

Various studies have previously reported the relevance of *B. cereus* spores in food poisoning caused by the consumption of contaminated raw food or vegetables [45, 46]. It is assumed that both vegetative cells and spores can reach the intestine, but the survival percentage of spores is considerably higher due to their resistance to stomach conditions [47, 48]. Therefore, based on this presumption, the few vegetative cells that survive are insufficient to reach the minimum infective dose required to cause disease (10^5^ – 10^8^ CFU) [12]. In keeping with this model, it is believed that diarrheic diseases provoked by enterotoxigenic *B. cereus* are associated with spore-contaminating foodstuffs, which germinate in the gut and produce the enterotoxins responsible for the symptoms [38, 48, 49]. In our study, we demonstrate that sporulation is not a determinant ecological trait for the persistence of enterotoxigenic *B. cereus* on ready-to-eat vegetables and that vegetative cells can reach the human guts and cause illness.

Sporulation and biofilm formation are two ecological traits contributing to the completion of the life cycle of *B. cereus*. Our genomic analysis demonstrated that strain DSM 2302, which is naturally unable to sporulate, lacks sporulation genes belonging to mid-late stages of sporulation. However, the first stages related to the phosphorelay and thus sporulation decision-making, as well as other cellular destinies, such as biofilms, are preserved [50, 51]. The inability to persist on/in melon leaves which was observed for both DSM 2302 or a *sigE* mutant of the emetic strain (AH187) that was affected in sporulation but not biofilm formation on melon leaves, indicated a major contribution of sporulation over biofilm formation in facilitating long-term bacterial survival in this ecological niche. Consistent with this, metabolomic analysis demonstrated that melon leaves are scarce in carbon and nitrogen sources and dominated by secondary metabolites, especially flavonoids, known as antibacterials and antibiofilms [52, 53, 54]. Contrary to this scenario, the endive is rich in nutrients and thus supportive of vegetative growth (**Fig. 3B**), most likely favouring biofilm formation as the preferred long-term bacterial survival strategy. We hypothesize that similar to the findings that have been reported for *B. subtilis* in the interaction with the rhizosphere, the collection of sensor kinases exposed on bacterial cell surfaces may discriminate the variety of signals and thus the final cellular decision-making [55]. Therefore, it remains to be determined how different metabolites found in our analysis are sensed by these *Bacillus* cells and integrated in the sporulation or biofilm pathways.

In addition to biofilm formation and sporulation, our genomic analysis showed a group of genes related to the use of plant carbon and nitrogen sources in the emetic strain AH187 (representative of *Bacillus* strain survival on plants) but not in DSM 2302. Even though the absence of these genes would be considered an adaptive disadvantage a priori, mutational analysis of these genes in strain AH187 did not indicate that they affect bacterial fitness on leaves. In alignment with our findings, a previous study also found that genes related to the use of plant carbon sources do not impart an ecological advantage; in fact, the presence of amino acids in food, such as vegetables, could improve bacterial adaptability [56]. However, considering that only viable single mutants could be tested, we could not determine whether this genetic background may collectively contribute to the survival of bacterial cells on plants, either as single vegetative cells or spores or as communities within biofilms. Overall, we conclude that adhesiveness of spores and their tolerance to external aggression can be overtaken by biofilm formation, and this switch is strongly influenced by the environment, where vegetative cells may persist on vegetables and reach humans.

Ii is generally assumed that in *B. cereus* life cycle spores reach the stomach and germinate, giving rise to metabolically active vegetative cells capable of colonizing the gut and producing the toxin responsible for damage to the host. We have, however, demonstrated that vegetative cells of DSM 2302 are able to overcome acid conditions typically found in the stomach after food ingestion (pH 4.5) (**Fig. 4A**) [20] for 5 h, which is sufficient time for the food to move forward from the stomach to the intestine [57, 58].

At this stage, two possible but non-exclusive bacterial behaviours are the formation of biofilms associated with the gut and the production of enterotoxins. Genomic analysis of DSM 2302 showed the presence of well-known genes of the biofilm pathway (regulatory and structural elements) and other genes whose products are related to such traits as bacterial cell adhesion to the host tissue, ECM assembly, swarming motility [59, 60, 61, 62], or bacterial invasion of epithelial cells, as previously reported for the human food poisoning pathogens *L. monocytogenes* and *E. coli* [43, 44]. In addition, DSM 2302 adheres to the bottom of the wells of static cultures, a low oxygen concentration environment that mimics the gut (**Fig. S4A**). DSM 2302 is genetically determined to produce the three primary known enterotoxins (Hbl, Nhe and Cytk) and the enterotoxins BceT, haemolysin I (cerolysin O), haemolysin II (HlyII), haemolysin III (HlyIII), and enterotoxin CwpFM (EntFM). BceT is lethal against mice, haemolysins are involved in lysing and permeabilizing host cells or intracellular organelles, and these toxins also have haemolytic activity against blood cells. EntFM is involved in bacterial virulence, motility, adhesion to epithelial cells and biofilm formation [63, 64, 65]. All these findings indicate that DSM 2302 has the necessary arsenal to adhere to epithelial cells of the gut and produce the toxins responsible for food poisoning in humans.

In summary, we identified a strain of *B. cereus* (DSM 2302) naturally impaired in sporulation but still able to complete a life cycle from vegetables to the final host (**Fig. 5**). Based on our findings, we propose that the persistence of this strain on plants relies on the metabolic nature of the host and the ability of the bacteria to form biofilms. Vegetative cells bearing structural elements related to biofilm formation (adhesins or EPSs) and protected by food can cross the stomach of humans and reach the intestine. The arsenal of biofilm-related structural factors would further ensure either the adhesion of cells to the intestine or the formation of biofilms, as well as the production of enterotoxins that ensure the release of nutrients from the human host. We propose that the loss of part of the sporulation programme and the reinforcement of bacterial traits related to adhesion, evasion of the immune system or production of toxins to scavenge nutrients are ecological traits of *B. cereus* fixed over evolution through pathogenic interactions with the human host. From a clinical perspective, our findings reinforce the presumed underestimation of food poisoning cases caused by *B. cereus* and the need to revise and improve the detection protocols that largely rely on the presence of spores or toxins in food [66, 67].

**Figure 5.**
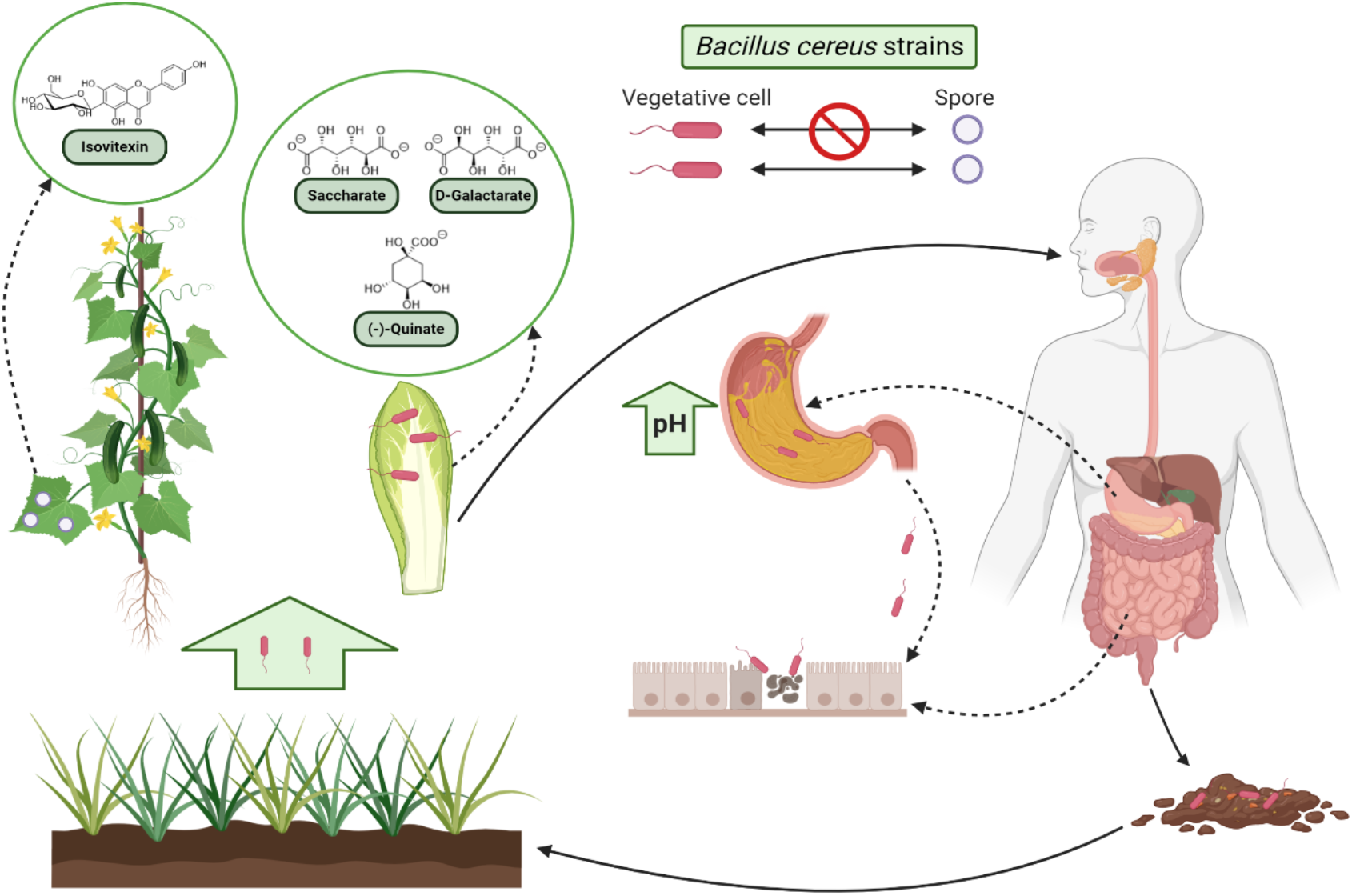
Reconsidering the life cycle of *B. cereus* from ready-to-eat vegetables to humans. Nonsporulating strains could survive over foodstuff products such as vegetables because of the large amount of nutrients compared to plant leaves, which are a more hostile niche. Furthermore, the ingestion of these contaminated foods increases stomach pH, and these conditions allow vegetative cells to survive stomach passage and enter the intestine to provoke human food poisoning.

## Acknowledgments

We would like to thank Josefa Gómez from the Ultrasequencing Unit of the SCBI-UMA for DNA sequencing, Zulema Udaondo from Bio-Iliberis and Rocío Bautista from Bioinformatic Unit of the SCBI-UMA for pangenome and genome analysis support, John Pearson from Bionand for guidance and assistance with microscopy images, María Isabel Castillo from Bionand-UMA for managing and feeding mice and Dr. Atul Parikh from UC Davis for access to his laboratory for the fabrication of PDMS leaf replicants used in this study. This work was supported by grants from European Research Council Starting Grant (BacBio 637971), and Plan Nacional de I+D+i of Ministerio de Economía y Competitividad, Ministerio de Ciencia e Innovación (AGL2016-78662-R and PID2019-107724GB-I00). Johan HL Leveau was financed by USDA-NIFA grant number 2014-67017-21695. Juan Antonio Guadix acknowledges financial support by grant (PIER-0084-2019) from Proyectos de Investigación en Salud de Junta de Andalucía.

