## Supplementary for "Sporulation is dispensable for the vegetable-associated life cycle of the human pathogen *Bacillus cereus*"

### Supplementary information

#### Supplementary Figures

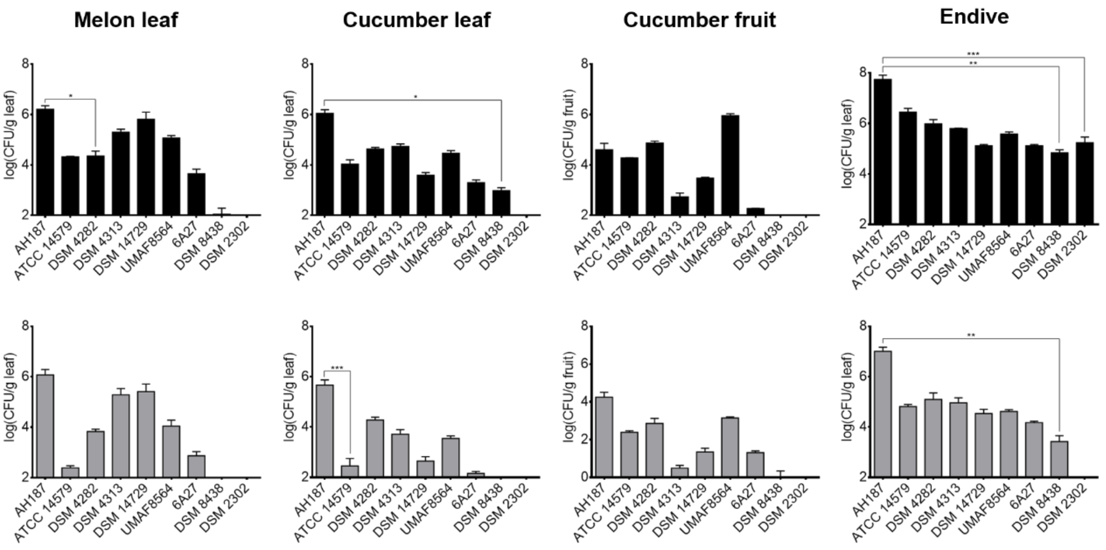


**Figure S1**. **Persistence and sporulation of *B. cereus* strains on melon and cucumber leaves, cucumber fruit or endive.** CFU counts per gram of leaf fruit 15 days after inoculation with *B. cereus* AH187, ATCC 14579, DSM 4282, DSM 4313, DSM 14729, UMAF8565, 6A27, DSM 8438 and DSM 2302. The top black bar plots show the counts of total CFUs over each surface. Bottom grey bars correspond to CFU counts of spores per gram of leaf or fruit. The detection limit of the technique is $10^2$ CFU/g of leaf. Error bars are the standard deviation of three or more replicates. Statistical analysis is the result of the comparison of all strains against AH187. * (P<0,05), ** (P<0,01), *** (P<0,001) (Dunn's test). No counts of the DSM 2302 strain appeared in any sporulation assay or on any surface, except endive. No significant counts of DSM 8438 strain appeared over all the surfaces (total CFUs and spores), except endive.

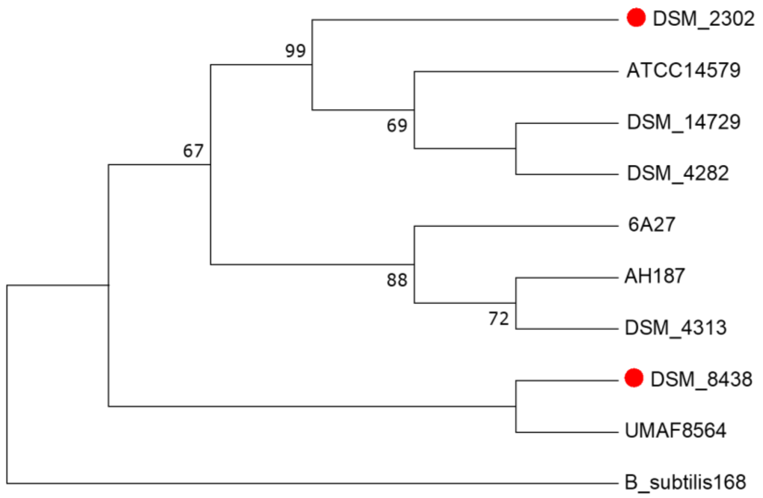


**Figure S2. Phylogenetic relation of *B. cereus* AH187, ATCC 14579, DSM 4282, DSM 4313, DSM 14729, UMAF8564, 6A27, DSM 8438 and DSM 2302 strains.** Red points indicate the strains that were not able to persist on the melon leaf surface (DSM 8438 and DSM 2302). *B. subtilis* subsp. *subtilis* 168 was used as an out-group. In the analysis, four concatenated genes (16S rRNA, *gyrB*, *rpoD* and *rpoB*) were handled to build a neighbour-joining tree using the MUSCLE algorithm of MEGA7.

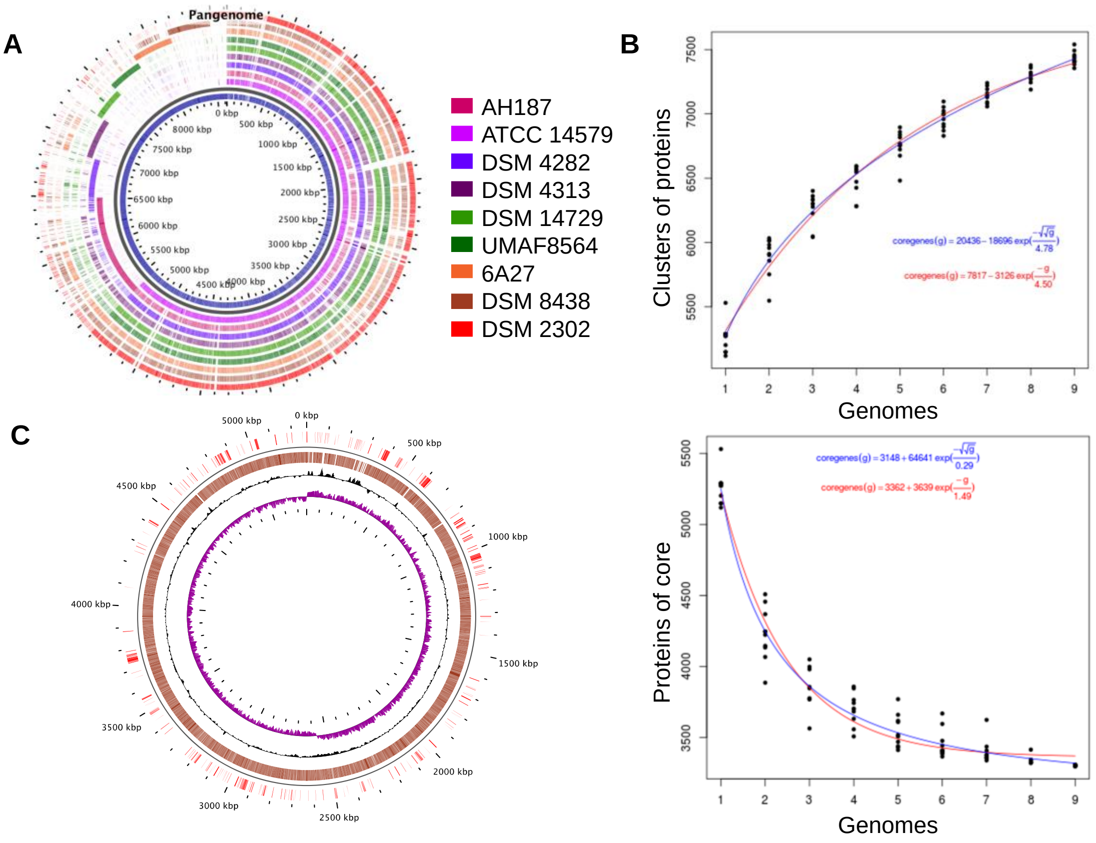


**Figure S3. Pangenome analysis of the strains AH187, ATCC 14579, DSM 4282, DSM 4313, DSM 14729, UMAF8564, 6A27, DSM 8438 and DSM 2302. A** Schematic view of the pangenome analysis of the nine studied strains. The inner layer represents the pangenome (all the analysed genes). Each colour layer corresponds to the genome of one B. cereus strain. **B** Statistical estimation with Willenbock (blue) and Tettelin (red) [72, 73]. The plot on the top points corresponds to pangenome size according to the number of genomes. In the plot on the bottom, the points correspond to the calculation of the core genome for each genome. **C** Schematic view of the genomic comparison between strains AH187 and DSM 2302. Genes present in strain AH187 are represented as brown lines throughout the genome, while genes absent in strain DSM 2302 are represented as red lines just above previous. The percentage of CG is represented in black, and the GC skew is represented in purple to locate the intergenic zones and the origins of replication of these strains, respectively.

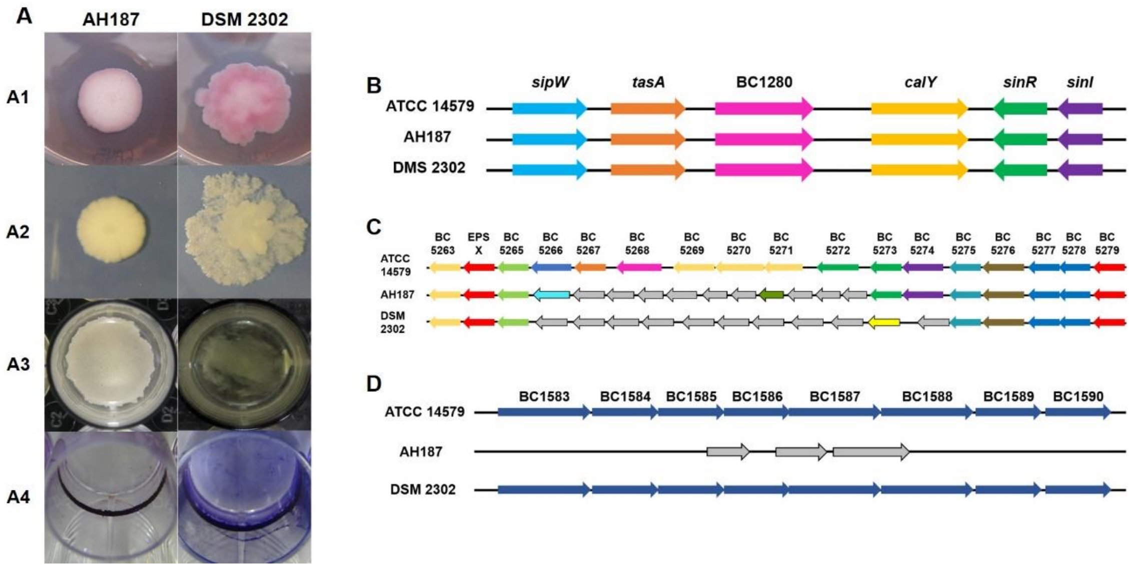


**Figure S4**. **Comparison of biofilm formation capacity of AH187 and DSM2302.** An *in vitro* biofilm and swarming assays of AH187 and DSM 2302 strains over 72 h on TY 1.5% agar supplemented with 20 µg/ml and 10 µg/ml Congo Red and Coomassie Brilliant Blue dyes, respectively (A1), TY 0.7% agar (A2) and TY broth (A3). A4 shows bacterial adhesion to an abiotic surface after staining with crystal violet. **B** *sipW-calY* locus implicated in extracellular matrix synthesis of AH187 and DSM 2302 compared to the *B. cereus* type strain (ATCC 14579). **C** Region homologous to the *eps1* region of ATCC 14579 involved in exopolysaccharide biosynthesis of AH187 and DSM 2302 strains. Middle grey genes correspond to different genes compared to each other. **D** Region homologous to the *eps2* region of ATCC 14579 involved in exopolysaccharide biosynthesis found in AH187 and DSM 2302 strains. Grey genes correspond to different genes located in the *eps2* genome region.

#### Supplementary Tables

**Table S1. Sequenced *B. cereus* strains deposited in Genbank.**

| Strain[a] | SUBID | BioProject | BioSample | Accession |
|---|---|---|---|---|
| AH187 | SUB3256016 | PRJNA17715 | SAMN02604058 | PHQS00000000 |
| ATCC 14579 | | | PRJNA384 | SAMN02603340 |
| DSM 4282 | SUB3254556 | PRJNA419655 | SAMN08094758 | PHKP00000000 |
| DSM 4313 | SUB3254567 | PRJNA419656 | SAMN08094759 | PHKQ00000000 |
| DSM 14729 | SUB3254577 | PRJNA419658 | SAMN08094761 | PHKS00000000 |
| UMAF8564 | SUB3254582 | PRJNA419659 | SAMN08094762 | PHKT00000000 |
| 6A27 | SUB3254521 | PRJNA419654 | SAMN08094757 | PHKO00000000 |
| DSM 8438 | SUB3254571 | PRJNA419657 | SAMN08094760 | PHKR00000000 |
| DSM 2302 | SUB3254588 | PRJNA224116 | SAMN03797561 | PHKU00000000 |

[a]*B. cereus* ATCC 14579, AH187 and DSM 2302 strains were previously deposited in Genbank.

**Table S2. Primers used to construct a *sigE* mutant of *B. cereus* AH187.**

| Primer name | Sequence |
|---|---|
| F_sigE_up/pMAD | cgatgcatgccatggtacccATGTTAGAGTTTGATACTACTCAAG |
| R_sigE_up | gtcttattctATTATCCCGCCTCCCATTATTAAG |
| F_sigE_down | gcgggataatAGAATAAGACCAGCCGTTATG |
| R_sigE_down/pMAD | cttctagaattcgagctcccTCACTTGCTTTCATCCTCC |

**Table S3**. **Pangenome analysis of *B. cereus* strains.** Core indicates the proteins shared among all the studied strains, accessory shows the proteins shared among all studied strains except one or less, and unique enumerates the proteins only present in one of the studied strains and not in the others. Differences in core proteins correspond to paralogous genes.

| ***B. cereus* strain** | **Core** | **Accessory** | **Unique** | **Total protein** |
|---|---|---|---|---|
| AH187 | 3424 | 1786 | 444 | **5654** |
| ATCC 14579 | 3413 | 1477 | 341 | **5231** |
| DSM 4282 | 3439 | 1989 | 391 | **5819** |
| DSM 4313 | 3420 | 1741 | 287 | **5448** |
| DSM 14729 | 3421 | 1692 | 248 | **5361** |
| UMAF8564 | 3412 | 1538 | 361 | **5311** |
| 6A27 | 3408 | 1687 | 280 | **5375** |
| DSM 8438 | 3406 | 1499 | 350 | **5255** |
| DSM 2302 | 3403 | 1573 | 414 | **5390** |

**Table S4**. **GO and KO terms, obtained from the pangenome analysis that are present in *B. cereus* DSM 2302 strains and absent in AH187 strain.** GO and terms associated with DSM 2302 strain with their corresponding function and DSM 2302 genes related with each GO/KO term.

| GO/KO | Function | GO/KO Term | Gene/s |
|---|---|---|---|
| GO:0004748 | Molecular Function | Ribonucleoside-diphosphate reductase activity, thioredoxin disulfide as acceptor | WP_001215825.1 |
| GO:0008374 | Molecular Function | O-acyltransferase activity | WP_061183899.1 |
| GO:0004668 | Molecular Function | Protein-arginine deiminase activity | WP_080444150.1 / WP_000435968.1 |
| GO:0009898 | Cellular Component | Cytoplasmic side of plasma membrane | WP_061182616.1 / WP_061182626.1 |
| GO:0031012 | Cellular Component | Extracellular matrix | WP_061183098.1 |
| GO:0019028 | Cellular Component | Viral capsid | WP_080444060.1 |
| GO:0000271 | Biological Process | Polysaccharide biosynthetic process | WP_016719053.1 |
| GO:0009446 | Biological Process | Putrescine biosynthetic process | WP_080444150.1 / WP_000435968.1 |
| K00558 | Cancer | DNA (cytosine-5)-methyltransferase 1 | WP_061182771.1 |
| K03196 | Infectious disease | Type IV secretion system protein VirB11 | WP_016718896.1 |
| K13735 | Infectious disease | Adhesin/invasin | WP_011040578.1 |
| K00463 | Infectious disease | Indoleamine 2,3-dioxygenase | WP_061183641.1 |
| K01467 | Drug resistance: antimicrobial | Beta-lactamase class C | WP_061182391.1 |
| K02172 | Drug resistance: antimicrobial | Bla regulator protein blaR1 | WP_061183726.1 |
| K05515 | Drug resistance: antimicrobial | Penicillin-binding protein 2 | WP_061182464.1 |
| K18143 | Drug resistance: antimicrobial | Two-component system, OmpR family, sensor histidine kinase AdeS | WP_061182681.1 |
| K00677 | Drug resistance: antimicrobial | UDP-N-acetylglucosamine acyltransferase | WP_062822245.1 |

| | | | |
|---|---|---|---|
| K03673 | Drug resistance: antimicrobial | Protein dithiol oxidoreductase (disulfide-forming) | WP_061183714.1 |
| K03760 | Drug resistance: antimicrobial | Lipid A ethanolaminephosphotransferase | WP_061183729.1 |
| K06079 | Drug resistance: antimicrobial | Copper homeostasis protein | WP_061182599.1 |

**Table S5. Genes absent in *B. cereus* DSM 2302 and DSM 8438.** Genomic comparison among persister and non persister strains reveal genes non present in DSM 8438 and DSM 2302 strains.

| **Gene/s** | **Annotation / Function** |
|---|---|
| BCAH187_A0689 | Response regulator. Two-component system CitB[b] |
| BCAH187_A2341 | Conserved hypothetical protein. ABC-2 type transport system permease protein[b] |
| BCAH187_A2533 | Transcriptional regulator, LysR family / Activator of inducible genes by hydrogen peroxide[a] |
| BCAH187_A2589 | Putative lipase. Acylhydrolase / Endoglucanase. Plant cell wall degradation[a] |
| BCAH187_A3065 | Hypothetical protein / Possible membrane protein[a] |
| BCAH187_A3159 | L-asparaginase ansA2 |
| BCAH187_A3164 | Pyrroline-5-carboxylase reductase |
| BCAH187_A3169 to BCAH187_A3171 | Two-component system and Glutaminase A[b] |
| BCAH187_A4674 | Cell surface protein |

**[a]**The annotation of the product corresponds to a possible function associated to amino acid sequence domains obtained by HHpred tool.
**[b]**The annotation of the product corresponds to GO/KO term or Pfam domain function.
